# Partial learning in human sound localization with asymmetric ears

**DOI:** 10.64898/2026.02.06.704341

**Authors:** Peter Bremen, Robert F. van der Willigen, A. John van Opstal, Marc M. van Wanrooij

## Abstract

Graphical abstract
Giving humans asymmetric barn-owl-like ears disrupts elevation localization by removing normal spectral pinna cues. With experience, listeners partially relearn to localize elevation using novel binaural cues. Results show that adult auditory spatial processing is flexible enough to repurpose cues, but strongly constrained in the extent of relearning.

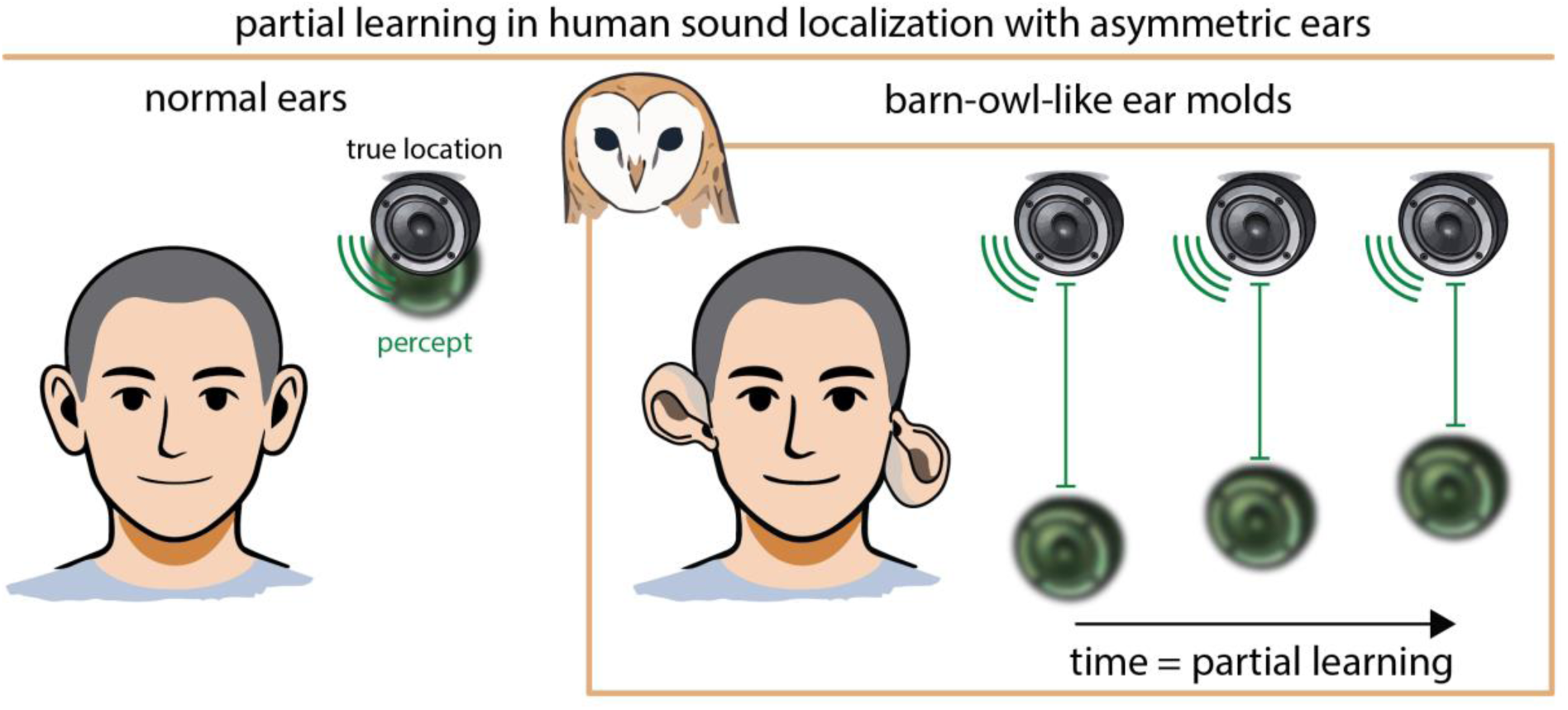

The brain computes sound location from auditory spatial cues. Humans and barn owls both localize sounds accurately, yet they rely on fundamentally different cue configurations shaped by their ear anatomy and neural circuitry. In humans, symmetrical ears provide interaural time and level differences (ITD and ILD) for horizontal localization, while vertical localization depends primarily on high-frequency, monaural spectral cues generated by the pinnae. Barn owls, by contrast, possess asymmetrical ears and use binaural cues for both azimuth and elevation. Because auditory pathways are tuned to species-specific cue statistics, it remains unclear whether humans can adapt when binaural level cues are also made informative about elevation. We tested this by fitting human listeners with asymmetric ear molds that disrupted normal spectral cues, introduced elevation-dependent ILDs, and left ITDs unaffected. Participants wore the molds during daily life and were tested with broadband, high-pass, and low-pass noise bursts. Acute mold exposure severely degraded elevation localization, whereas horizontal localization remained largely unaffected. With prolonged exposure, elevation localization improved, but adaptation was limited, variable across listeners, and fluctuated across sessions. Improvement was strongest for broadband sounds. Because broadband and high-pass sounds both contained high-frequency information, this broadband advantage cannot be explained by access to additional spectral cues alone. Instead, it suggests that low-frequency ITDs helped listeners exploit the altered binaural cue structure, including the elevation-dependent ILDs introduced by the molds. These findings show that humans can partially adapt to asymmetric outer-ear acoustics when complementary spatial cues help resolve ambiguity in the altered cue structure.

## Introduction

Both humans and barn owls excel at localizing sounds, despite having vastly different auditory anatomies and neural architectures. Barn owls possess asymmetrically placed ears (Knudsen and Konishi, 1979; Coles and Guppy, 1988; von Campenhausen and Wagner, 2006) and specialized neural mechanisms, including a topographic auditory space map in the midbrain (Knudsen and Konishi, 1978), which are crucial for their exceptional localization abilities (Payne, 1971; Konishi, 1971). In contrast, humans have symmetrical ears (Strutt, 1907; Wightman and Kistler, 1989; Middlebrooks and Green, 1991; Blauert, 1997; Van Wanrooij and Van Opstal, 2005) and do not appear to possess a topographic auditory space map in the ascending auditory pathway (Zwiers et al., 2004; Middlebrooks, 2021), yet they localize sounds with comparable accuracy (Middlebrooks and Green, 1991; Heffner and Heffner, 1992; Frens and Van Opstal, 1995). Across the lifespan, humans encounter changes in ear shape and hearing sensitivity that alter spatial cues. The human auditory system shows remarkable plasticity in adjusting to such changes throughout life (Otte et al., 2013), including through learning after wearing ear molds that disrupt pinna cues (Hofman et al., 1998; Van Wanrooij and Van Opstal, 2005; Carlile, 2014), following acute unilateral hearing loss by plugging the ear (Kumpik et al., 2010; Kumpik and King, 2018), adjusting to non-individualized or supernormal HRTFs (Shinn-Cunningham et al., 1998), and spectrally warped or band-limited spatial information (Majdak et al., 2013). Visual input can further support auditory spatial recalibration (Radeau and Bertelson, 1974; Zwiers et al., 2003; Recanzone, 1998; Zonooz and Van Opstal, 2019; Kayser et al., 2023). However, there are limits to this plasticity: severe hearing loss or extreme manipulations such as swapping ear inputs can prevent adaptation (Hofman et al., 2002; Van Wanrooij and Van Opstal, 2004, 2005; Otte et al., 2013). Here, we investigated how humans adapt when challenged with asymmetric outer-ear acoustics that introduce elevation-dependent binaural level cues.

In humans, the brain relies on three main acoustic cues to determine sound direction: interaural time differences (ITDs, Fig. 1a) and interaural level differences (ILDs, Fig. 1b) provide horizontal (azimuth) information, while monaural spectral shape cues (Fig. 1c), created by the pinnae, provide vertical (elevation) information (Strutt, 1907; Middlebrooks and Green, 1991). These cues are bandwidth-dependent: ITDs are most effective at low frequencies (<1500 Hz), ILDs at high frequencies (>3000 Hz), and spectral cues above ∼4000 Hz (Middlebrooks and Green, 1991; Wightman and Kistler, 1992; Macpherson and Middlebrooks, 2002). These cues are processed along dedicated brainstem pathways and contribute to a perceptual representation of sound space (Yin et al., 2019). Barn owls, due to their smaller heads, use ITDs at higher frequencies than humans (Fig. 1d). For elevation, they do not rely on spectral cues (Fig. 1f), but on elevation-dependent ILDs (Fig. 1e) arising from asymmetrically positioned and oriented ears (Knudsen and Konishi, 1979; Coles and Guppy, 1988; von Campenhausen and Wagner, 2006). Thus, their spatial hearing system differs fundamentally from the human system in both peripheral anatomy and central processing (Knudsen and Konishi, 1978; Middlebrooks, 2021).

**Figure 1.**
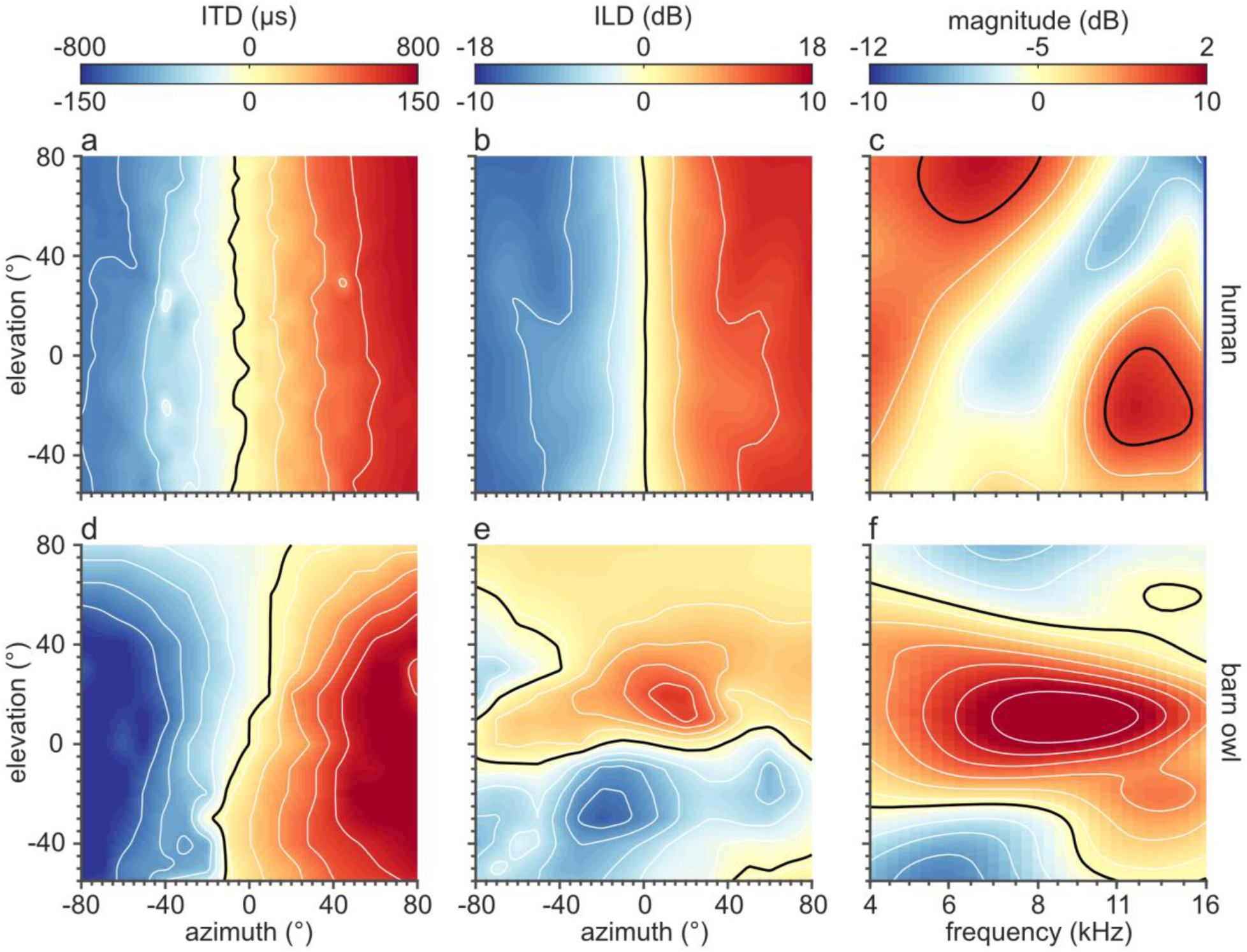
Acoustic spatial cues in humans and barn owls. Heat maps show the distribution of three key spatial hearing cues: interaural time differences (ITDs; a,d), interaural level differences (ILDs; b,e), and monaural spectral shape cues at 0° azimuth (c,f), for humans (top row) and barn owls (bottom row). The black line in each panel indicates the zero (dB, or µs) contour line. In both species, ITDs vary systematically with azimuth (a,d). ILDs vary primarily with azimuth in humans (b), but with elevation in barn owls (e), due to their asymmetrical ear placement. Spectral shape cues in humans (c) exhibit a distinct elevation-dependent notch between 4 and 16 kHz, which is absent in barn owls (f). Color bars above each column indicate the value scales for humans (top labels) and barn owls (bottom labels). ITDs were extracted from low-pass filtered directional transfer functions (DTFs) below 1 kHz for humans and below 16 kHz for barn owls. ILDs were calculated over 4-16 kHz (humans) and 1-12 kHz (barn owls). Spectra in panels (c) and (f) were smoothed using a Gaussian filter (Q=8) to highlight perceptually relevant peaks and notches. Data adapted from Bremen et al. (2010) for humans (participant 2 in the current study) and Bremen et al. (2007) for barn owls.

We therefore asked whether humans can adapt to a radically different cue configuration inspired by one key feature of barn-owl spatial hearing: asymmetric, elevation-dependent binaural level cues. The barn owl comparison served as biological motivation rather than as an attempt to reproduce barn-owl acoustics or neural cue processing. Using custom-made asymmetrical ear molds, we introduced combined azimuth and elevation-dependent ILDs in human listeners, reduced spectral shape cues, and left ITDs largely unaltered. Participants wore the molds during everyday life for up to five weeks, and we assessed localization without visual feedback or laboratory training using broadband, high-pass, and low-pass noise bursts. This allowed us to test whether auditory spatial plasticity is limited to recalibration within established cue-to-space mappings, or whether listeners can partially adapt binaural cues across spatial dimensions when the acoustic cue structure is altered.

## Methods

### Ethics

All experiments were conducted in 2008 and 2010 and complied with institutional and international ethical standards for research involving human participants. All participants gave informed written consent after being fully informed of the procedures. The experimental protocol was approved by the Local Ethics Committee of Radboud University Nijmegen and adhered to the principles of the Declaration of Helsinki (2008).

### Participants

Eleven participants (ages 23-52; 4 females), including three authors (P1-3), took part in the study, split in two cohorts (Table 1). In the 2008 cohort, P1–P3 were authors and P4 was a PhD student in the research group. In the 2010 cohort, all participants other than P1 were paid volunteers recruited externally from the Radboud University student population. All had normal hearing, as confirmed by pure-tone audiometry showing thresholds within 20 dB across frequencies from 0.25 to 11.3 kHz in eleven 0.5-octave steps. All participants had normal or corrected-to-normal vision, except for Participant 2, who did not wear his prescription glasses during the experiment, and Participant 3, who was amblyopic in the right eye.

**Table 1.** Overview of the experimental paradigm per participant, summarizing to which cohort each participant belonged, and how many days they wore the molds.

| Participant | Cohort | Period (days) |
| --- | --- | --- |
| 1 | 2008 | 9 |
| 2 | 2008 | 48 |
| 3 | 2008 | 59 |
| 4 | 2008 | 42 |
| 1 | 2010 | 19 |
| 5 | 2010 | 7 |
| 6 | 2010 | 5 |
| 7 | 2010 | 9 |
| 8 | 2010 | 9 |
| 9 | 2010 | 3 |
| 10 | 2010 | 27 |
| 11 | 2010 | 3 |

### Setup

The experimental setup has been described in detail previously (Bremen et al., 2010). Participants sat in the center of a motorized vertical loudspeaker hoop (2.5 m diameter) mounted in a sound-attenuated, dark room (3 × 3 × 3 m) lined with acoustic foam (50 mm base, 30 mm pyramids; Uxem Flexible Foams, Lelystad, NL). Fifty-eight loudspeakers (Visaton SC 5.9, art. no. 8006) were mounted on the hoop, allowing sound presentation from various spatial locations defined in double-polar coordinates (Knudsen and Konishi, 1979), with azimuth (horizontal angle) and elevation (vertical angle) specified in degrees.

A PC (2.8 GHz Intel Pentium D; Dell) controlled speaker selection, sound presentation (RP2.1; Tucker-Davis Technologies, 48,828 Hz), head movement tracking (RA16; Tucker-Davis Technologies, 1017 Hz), and directional transfer function measurements. We recorded head movements using a magnetic search-coil technique (Robinson, 1963). Participants wore a lightweight glasses frame (without lenses) fitted with a search coil and a red laser pointer (LQB-1-650; World Star Tech) aimed at a 1-cm disk positioned 30 cm in front of the eyes. This ensured visibility of the pointer while preventing distracting reflections from walls or speakers. We converted coil voltages to degrees of rotation using calibration procedures described by Goossens and Van Opstal (1997).

### Sounds

For behavioral testing, we used Gaussian white-noise (GWN) bursts filtered into three spectral bands: broadband (BB, 0.5-20 kHz), high-pass (HP, 3-20 kHz), and low-pass (LP, 0.5-1.5 kHz). These passbands were chosen to emphasize different spatial cue combinations but should not be interpreted as perfectly isolating single cues. Broadband stimuli provided the richest cue set, including low-frequency ITDs, high-frequency ILDs, and high-frequency monaural spectral cues. High-pass stimuli emphasized high-frequency ILDs and pinna-based spectral structure, while excluding the low-frequency fine-structure ITDs that normally dominate human azimuth localization. Although envelope ITDs may in principle be present in high-frequency noise bursts, they were not expected to provide a dominant cue in these short broadband-noise stimuli, especially in the presence of stronger high-frequency level and spectral cues. Low-pass stimuli emphasized low-frequency ITDs and largely removed high-frequency pinna-based spectral cues. Any elevation-dependent level cues from torso or head filtering in this low-frequency range were expected to be weak and unreliable, particularly because the LP band ended at 1.5 kHz. Thus, the cue labels BB, HP, and LP refer to the dominant cue information available in each stimulus condition rather than to complete cue isolation.

Each burst lasted 150 ms, including 5-ms sine²/cosine² onset and offset ramps. We generated 100 unique noise realizations per spectral condition and presented them in randomized order during testing sessions. To prevent participants from using absolute intensity as a localization cue, we randomly varied the sound level. For the 2008 cohort, the sound levels varied between 40 and 70 dBA in 10-dB steps. Some sessions included only BB sounds at 50 and 70 dBA. For the 2010 cohort, levels were 50, 60, or 70 dBA. Sound levels were measured at the participant’s head position using a BK2610 sound level meter and BK4144 microphone (Brüel and Kjær, Nærum, Denmark). All levels were clearly audible and within a moderate range; level-dependent localization was therefore not tested as a separate experimental factor.

### Ear Molds

We constructed custom ear molds for each participant by filling the concha with silicone casting material (Otoform Otoplastik-K/c; Dreve, Unna, Germany), leaving the ear canal unobstructed to maintain acoustic access. Inspired by the asymmetrical ear morphology of barn owls, we shaped the molds to introduce directional reflectors: the left mold featured a flap above the ear canal to enhance sensitivity to sounds from below, while the right mold had a flap below the ear canal to enhance sensitivity to sounds from above (Fig. 2). Note that the molds were individually and manually shaped for each participant to produce the same qualitative asymmetry, but the fabrication procedure was not standardized to impose identical acoustic transformations across listeners.

**Figure 2.**
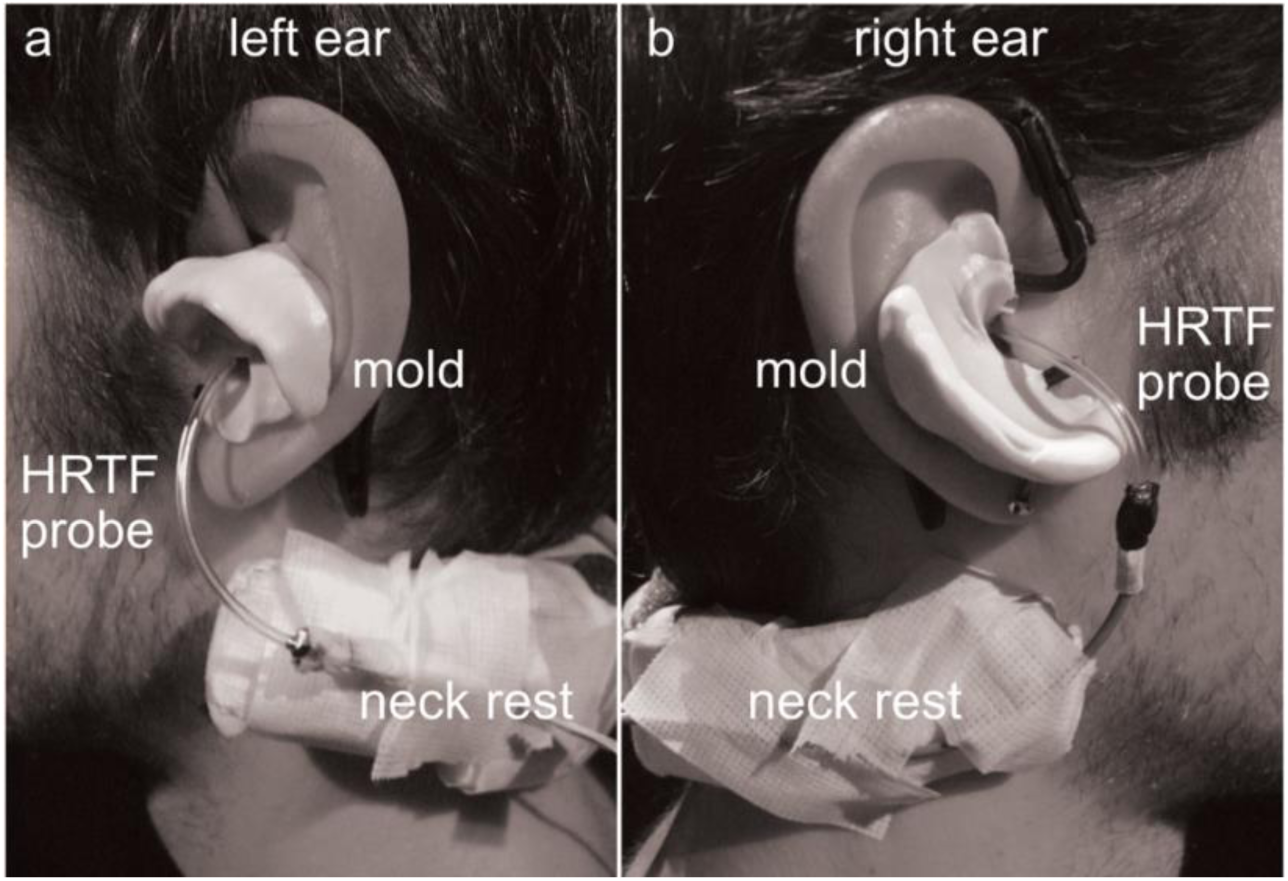
Participant 2 wearing asymmetric ear molds. Side views of the participant’s left ear (a; flap above ear canal) and right ear (b; flap below the ear canal). Ear hooks that support the molds and microphones for measuring head-related transfer functions (HRTFs) can also be seen. Note that the neck rest was used to keep the head still during HRTF measurements, but was removed for the sound-localization measurements

To characterize the acoustic transformations produced by the molds, we analyzed HRTFs measured in one participant (P2) with and without the molds, following Bremen et al. (2010). HRTFs were also measured in additional participants at the time of testing, and these measurements were used to verify that the molds disrupted the normal spatial cues and produced cue transformations qualitatively similar to those observed for P2. However, because the raw HRTF data from these additional participants are no longer available for reanalysis, the acoustic analyses presented here are restricted to P2 and serve as an example of the cue transformations produced by the molds.

Briefly, for visualization and analysis, the resulting magnitude spectra were converted to sound level (dB) and smoothed using a Gaussian filter with a constant Q-factor of 8 (Algazi et al., 2001; Bremen et al., 2010). This procedure removes high-frequency fluctuations to which the human auditory system is relatively insensitive while preserving the primary direction-dependent spectral features. With the molds in place, interaural time differences (ITDs) remained unaffected, varying systematically with azimuth (Fig. 3a; cf. Fig. 1a). In contrast, interaural level differences (ILDs) changed markedly, varying not only with azimuth but also with elevation (Fig. 3b; cf. Fig. 1b). This elevation dependence can be seen from the tilted iso-ILD contours: at a given azimuth, ILDs became more positive with increasing elevation. The molds therefore introduced an elevation-dependent component into the ILD cue. To quantify the relative elevation dependence of the ILD cue, we fitted a plane to the ILD map as a function of azimuth and elevation and compared the elevation and azimuth coefficients. Without molds, human ILDs were almost exclusively azimuth-dependent, with an elevation-to-azimuth gradient ratio of 0.00. With the molds, this ratio increased to 0.21, indicating that the molds introduced a measurable elevation-dependent component into the ILD cue. This component remained much weaker than in the barn owl, for which the elevation-to-azimuth ratio was 1.95, consistent with ILDs being dominated by elevation rather than azimuth. Thus, the molds did not reproduce barn-owl acoustics, but introduced a weak, systematic elevation-dependent ILD component into the human cue set.

**Figure 3.**
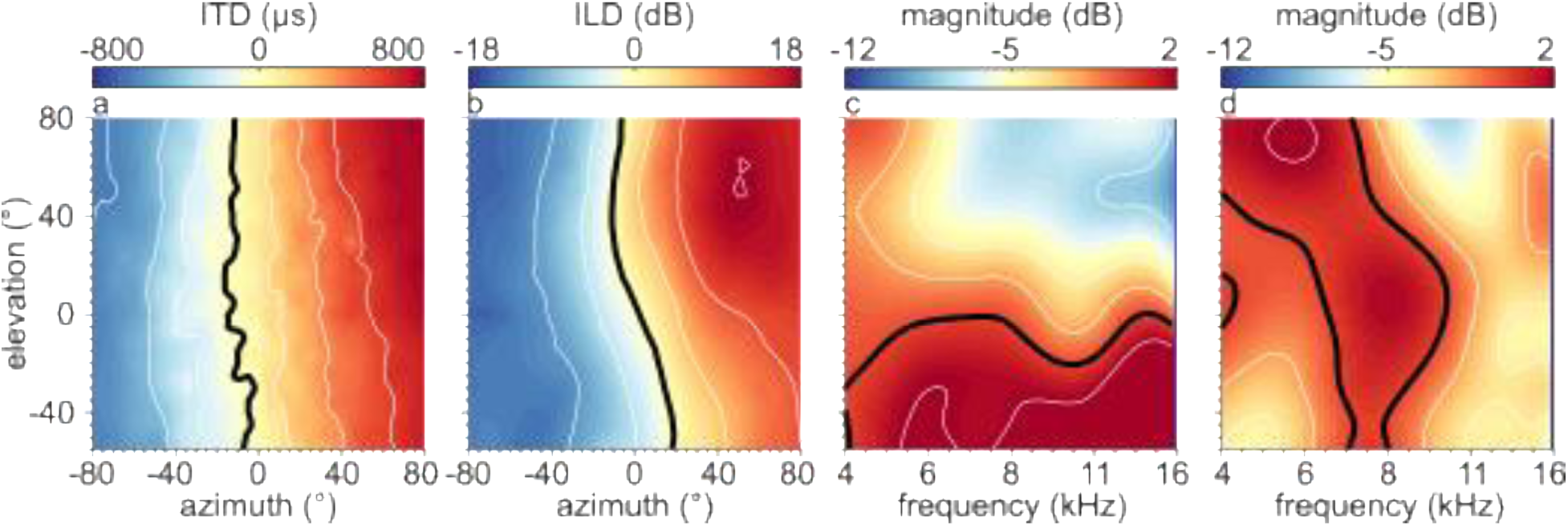
Binaural and monaural cues for participant 2 wearing barn-owl molds. (a) Interaural time differences (ITDs) vary systematically with azimuth. (b) Interaural level differences (ILDs) vary with both azimuth and elevation. The elevation dependence is visible as tilted iso-ILD contours: for many azimuths, increasing elevation shifts the ILD toward more positive values. (c) Spectral shape cues in the midsagittal plane for the left ear show attenuation for sounds from above between 4 and 16 kHz, consistent with the mold’s design. (d) The right ear mold did not produce a clear elevation-dependent spectral pattern. The black line in each panel indicates the zero-contour line.

Monaural spectral shape cues were significantly disrupted (Fig. 3c), with the elevation-dependent spectral notch, seen under normal conditions (Fig. 1c), largely eliminated. Instead, the left ear exhibited a broad, frequency-independent change: attenuation of approximately −8 dB for sounds from above and an increase of around +5 dB for sounds from below. The right ear showed no consistent elevation-related pattern (Fig. 3d).

In summary, the molds preserved ITDs as cues for azimuth (Fig. 3a), but 1) disrupted reliable monaural spectral cues (Fig. 3c,d), and 2) introduced elevation-dependent ILDs (Fig. 3b). Unlike in barn owls, however, ILDs with the molds continued to vary with azimuth, meaning that human listeners would have to disambiguate azimuthal and elevational contributions to ILDs. This is an additional challenge that barn owls, with their fixed spatial mapping of ILDs, do not face (cf. Fig. 1e).

### Behavioural Testing

Each experimental session consisted of a series of open-loop localization trials. At the start of each trial, a central fixation LED turned on. Participants aligned the head-fixed laser pointer with the LED and initiated the trial by pressing a button. After a random delay (300-1000 ms in 100-ms steps), the fixation light turned off and a sound was presented from one of the loudspeakers. Participants were instructed to point the head-fixed laser “as quickly and as accurately as possible” toward the perceived sound location. Reaction times exceeded the stimulus duration by more than 150 ms, ensuring open-loop conditions (i.e., responses were not guided by ongoing sound). Sessions typically lasted under an hour and were divided into several blocks with short (5-minute) breaks between them, in which the room lights were turned on.

The range of sound-source locations differed between the two participant cohorts. The 2008 cohort was tested with a large spatial range: −75° to +75° in azimuth, and −57.5° to +75° in elevation. The 2010 cohort was tested within a narrower spatial range: −35° to +35° in both azimuth and elevation.

### Adaptation Period

Participants wore their custom-made, barn-owl molds for durations ranging from 3 to 59 days (Table 1). Participant 1 participated in two separate adaptation periods, once in 2008 and once in 2010, with different molds in each case. Participants wore the molds throughout the day during their normal daily activities and during all behavioral testing sessions, removing them only during sleep.

Sound-localization behavior was assessed frequently: before mold insertion and on nearly every weekday throughout the adaptation period. Importantly, no performance feedback was provided during testing sessions. Participants received only incidental feedback from their daily interactions with the environment.

## Data Analysis

### Saccade Detection

We processed and analyzed all behavioral data using MATLAB (The MathWorks, Natick, MA, USA). Head saccades, defined here as rapid goal-directed head-orienting movements toward the perceived sound location, were detected automatically. Movement onset and offset were identified using velocity thresholds of 20°/s and 15°/s, respectively. All detections were visually inspected, and corrections were made manually when necessary.

### Localization Performance

We quantified sound localization performance using linear regression:

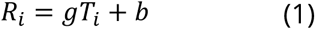

where *T*_i_ and *R*_i_ denote the target and response locations, respectively, for trial i and pooled across sound levels. The slope g (dimensionless) reflects the gain, indicating the strength of the stimulus-response relationship, and the intercept b (in degrees) represents the bias in responses. Perfect localization corresponds to a gain of 1 and a bias of 0°.

We computed gains and biases separately for azimuth and elevation, for each participant, session, and sound type. The gain is our primary measure of localization performance, since biases showed no systematic variation across sessions. We also calculated localization errors, which gave similar results as the gain (not shown).

In addition to the primary regressions, we also regressed response azimuth on stimulus elevation to quantify potential cross-dimensional effects. This analysis tested whether the altered cues, specifically, the introduction of elevation-dependent ILDs, induced systematic changes in azimuth responses as a function of elevation.

### Statistics

We used estimation statistics to quantify differences between experimental conditions (Calin-Jageman and Cumming, 2019; Ho et al., 2019). We obtained 95% confidence intervals (CI) of the mean difference distributions by performing a bootstrap analysis with replacement. Results are reported as the mean and the 95% confidence interval (CI) of the gain, in the format: mean [lower CI, upper CI]. Additionally, we report p-values from two-sided permutation t-tests, which indicate the probability of observing an effect at least as extreme as the measured effect under the null hypothesis of no difference.

## Results

### Impact of Molds - Example Participant

To illustrate the effects of the ear molds and our linear-regression-based analysis approach, Figure 4 presents stimulus-response data from one example participant (P10), tested at three key stages: before mold insertion (yellow), immediately after insertion (blue), and after 27 days of continuous use (red). This example demonstrates how an abrupt change in spatial cues affects sound localization and how performance evolves with prolonged exposure. Also, it provides a visual check that the linear regression gain captures the main structure of the localization responses and highlights the acute effects, partial recovery, and cross-dimensional response patterns that motivated the subsequent group-level analyses.

**Figure 4.**
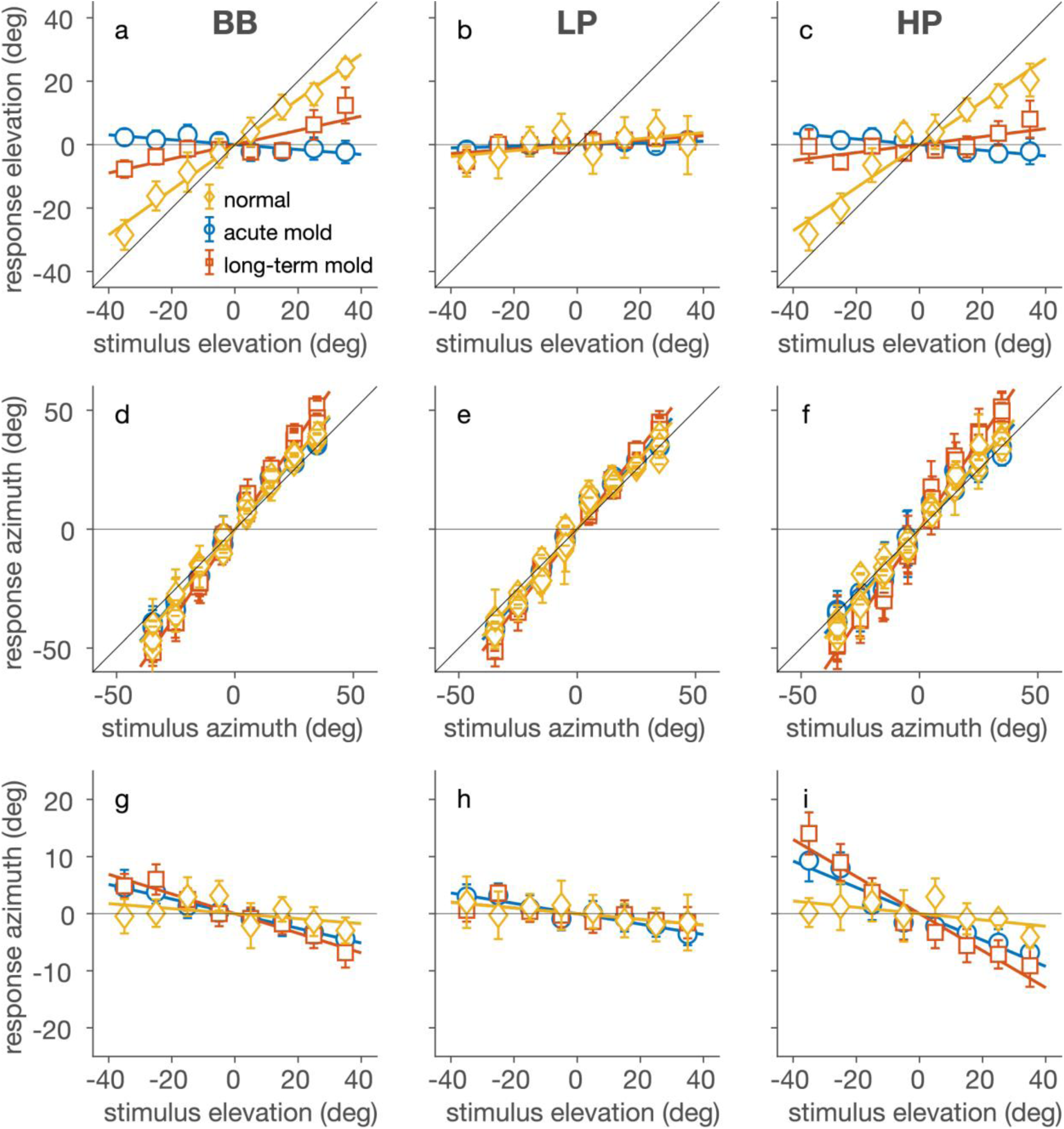
Example sound localization before and after exposure to barn-owl molds. Stimulus-response plots for participant 10 show sound localization behavior in elevation (a-c) and azimuth (d-f) across three conditions: without molds (yellow), immediately after mold insertion (blue), and after 27 days of mold use (red). Panels g-i show cross-dimensional effects, plotting response azimuth as a function of stimulus elevation. Each column corresponds to a different sound type: broadband (BB, left), high-pass (HP, middle), and low-pass (LP, right) noise. Markers indicate the mean response, and error bars show 95% confidence intervals. Lines represent linear regression fits. Data were pooled across sound levels and across the orthogonal target dimension. This example is shown to illustrate the stimulus response patterns and regression-based analysis, not to support group-level conclusions. Group-level effects and individual variability are quantified in Figs. 5–7.

Figure 4 depicts stimulus-response relationships for elevation (Fig. 4a-c) and azimuth (Fig. 4d-f), as well as cross-dimensional effects, showing how azimuth responses depend on stimulus elevation (Fig. 4g-i). Results are shown separately for broadband, high-pass, and low-pass sounds. Data were pooled across sound levels and across the orthogonal spatial dimension. Thus, elevation-response plots were pooled across target azimuths, azimuth-response plots across target elevations, and azimuth-versus-elevation plots across target azimuths. Before wearing the molds, the participant localized broadband and high-pass sounds accurately in elevation (Fig. 4a,c, yellow; gains ≈ 0.7), but performed poorly for low-pass sounds (Fig. 4b, yellow; gain ≈ 0), consistent with the absence of high-frequency spectral cues in low-pass stimuli. Immediately after mold insertion, elevation localization for broadband and high-pass sounds collapsed (Fig. 4a,c, blue; gains near zero), while low-pass performance remained unchanged and poor (Fig. 4b, blue). This acute deficit confirms that the molds effectively disrupted the spectral cues normally used for sound elevation localization.

After 27 days of wearing the molds, elevation performance partially recovered. Gains increased to approximately 0.3 for broadband sounds and 0.2 for high-pass sounds (Fig. 4a,c, red), indicating that the participant began to extract useful elevation information from the altered acoustic cues. As expected, given the lack of informative spectral structure, low-pass elevation localization remained poor throughout (Fig. 4b, red).

In contrast, azimuth localization remained robust across all conditions (Fig. 4d-f), with gains close to 1.0 before mold insertion (yellow), immediately after insertion (blue), and after prolonged use (red). This stability is consistent with the preservation of low-frequency ITDs and reliable ILDs for horizontal localization. However, the molds also introduced a systematic cross-dimensional effect: azimuth responses became dependent on stimulus elevation (Fig. 4g-i). Specifically, sounds presented at higher elevations tended to be localized more toward the left. This effect was most pronounced for high-pass sounds, for which the azimuth-versus-elevation regression gain increased to about 0.2 after adaptation (Fig. 4i, red), suggesting that the elevation-dependent ILDs introduced by the molds began to influence azimuth judgments. For the BB sounds, the complementary use of unaffected ITD cues decreased this effect considerably.

In summary, this example participant showed a pronounced and immediate loss of elevation localization after mold insertion, accompanied by the emergence of cross-dimensional interference in azimuth. With extended exposure, elevation gains partially recovered, demonstrating experience-dependent adaptation to the altered acoustic cues. Note that across all panels, the localization gain (the slope of the stimulus-response regression, Eq. 1) provides a compact and informative summary of localization behavior. In the following sections, we show that these acute effects and gradual adaptations generalize across participants, as observed in their localization gains over time.

### Acute Impact of Molds

We quantified the acute effects of the asymmetric ear molds by comparing sound-localization performance before and immediately after mold insertion across all participants (Fig. 5). We assessed changes in localization gain (Eq. 1) along three dimensions: elevation (Fig. 5a,d), azimuth (Fig. 5b,e), and the dependence of azimuth responses on stimulus elevation (Fig. 5c,f). Each analysis was performed separately for broadband (BB; blue), low-pass (LP; red), and high-pass (HP; yellow) noise bursts, allowing us to isolate how abrupt cue perturbations affect different components of spatial hearing.

**Figure 5.**
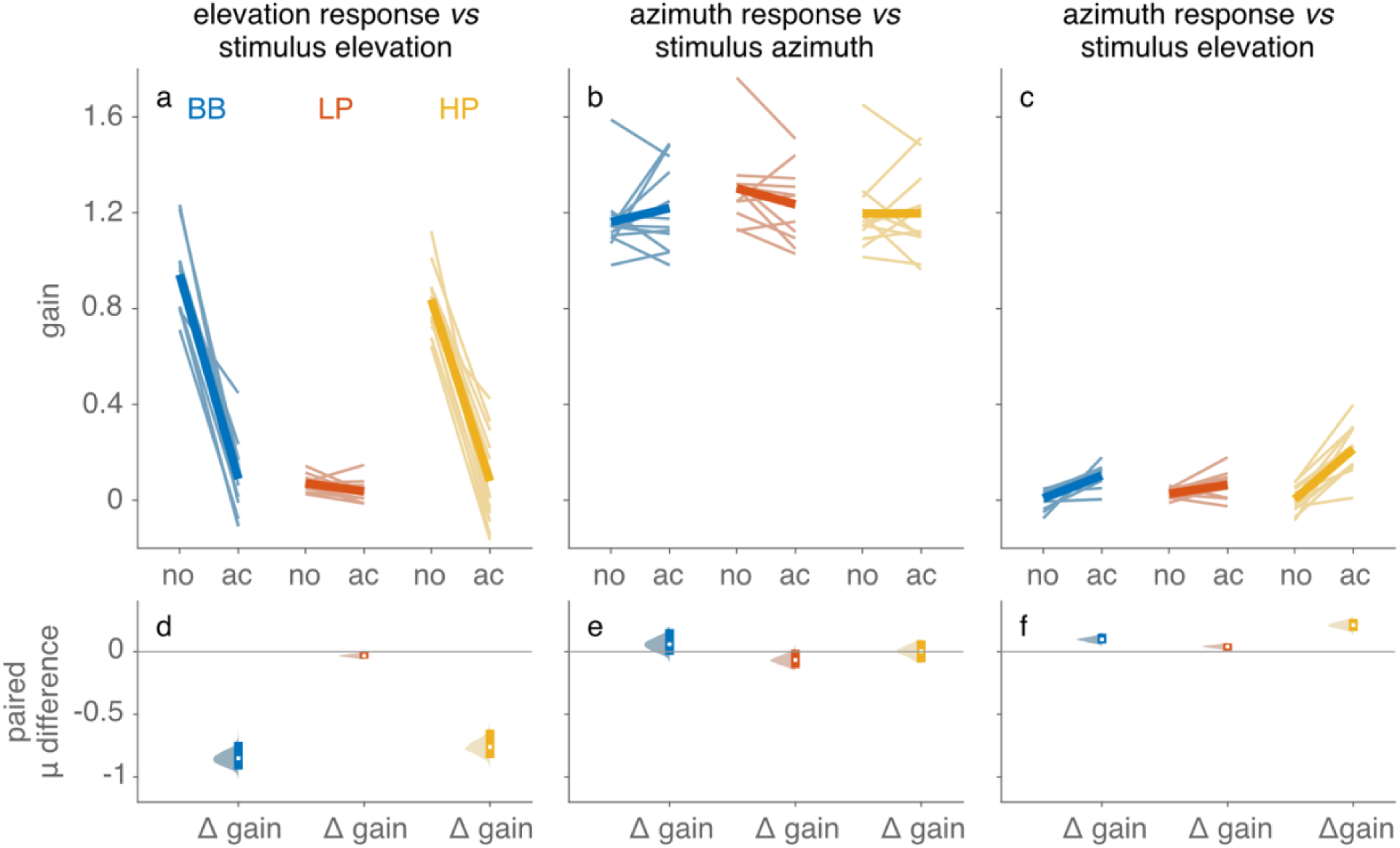
Acute effects of asymmetric ear molds on human sound localization performance. Sound localization gains are shown for elevation (a, d), azimuth (b, e), and the interaction between azimuth responses and stimulus elevation (c, f), for three spectral noise conditions: broadband (BB; blue), low-pass (LP; red), and high-pass (HP; yellow). Each plot compares performance without molds (“no”) and with acutely worn molds (“ac”). Each panel in the top row (a-c) displays slope graphs where individual paired observations (each participant’s performance without vs. with molds) are connected by thin lines and the group averages (thick lines). Panels d-f present the corresponding paired differences (acutely worn - no mold) between conditions where the white marker represents the mean difference, the vertical error bar marks the 95% confidence interval, and the colored patch illustrates the bootstrap sampling distribution.

Participants localized broadband and high-pass sounds accurately in elevation before wearing the molds (Fig. 5a; mean gain BB = 0.94 [0.87, 1.04], where square brackets indicate bootstrapped 95% confidence intervals; gain HP = 0.84 [0.77,0.92]). Immediately after mold insertion, the gains dropped by ΔBB = −0.85 [0.94, −0.71] (p<0.001) and ΔHP = −0.76 [−0.85, − 0.64] (p<0.001) (Fig. 5d), indicating a near-complete loss of vertical localization. This dramatic decline confirms that the molds effectively disrupted the spectral cues required for elevation sound localization. In contrast, low-pass sounds were localized poorly in elevation both before (LP gain = 0.07 [0.06, 0.09]) and after mold insertion (LP gain = −0.04 [0.02, 0.07]), consistent with the absence of informative high-frequency spectral structure. Comparing gains directly (Fig. 5d) shows that the acute loss of elevation performance occurred selectively for broadband and high-pass stimuli.

By contrast, azimuth localization remained robust across all conditions. Gains were close to 1.2 [1.1, 1.3] both before and after mold insertion, with no reliable change induced by the molds (Δ_BB_: 0.06 [−0.04, 0.16]; *p* = 0.31; Δ_LP_: −0.07 [−0.14, 0]; *p* = 0.10; Δ_HP_: 0 [−0.09, 0.08]; *p* = 0.98; Fig. 5b,e). This stability is consistent with the preservation of low-frequency ITDs and reliable ILDs, which continued to provide valid horizontal localization cues.

A systematic effect emerged, however, when we examined how azimuth responses depended on stimulus elevation (Fig. 5c,f). This interaction was negligible at baseline but increased significantly after mold insertion, indicating cross-dimensional interference. The azimuth-elevation coupling increased for all sound types, with the strongest effect observed for high-pass sounds (Δ_HP_: 0.21 [0.16, 0.27]; *p* ≪ 0.01), followed by broadband sounds (Δ_BB_: 0.1 [0.06, 0.14]; *p* ≪ 0.01). Low-pass sounds showed a smaller, yet reliable, increase (Δ_LP_: 0.04 [0.01, 0.06]; *p* = 0.03). These results indicate that the molds introduced elevation-dependent structure into binaural cues that are normally interpreted purely in terms of azimuth.

In summary, acute exposure to the asymmetric molds caused a severe disruption of sound elevation localization, selectively affecting sounds that normally rely on high-frequency spectral cues, while horizontal localization remained largely intact. At the same time, the emergence of elevation-dependent biases in azimuth responses shows that the altered binaural cues were interpreted according to the normal cue-to-space mapping: changes in ILD were treated as changes in azimuth, even though the molds had made ILDs partly dependent on elevation.

### Time course of Adaptation

To visualize the dynamics of adaptation, we plotted elevation gains for each participant as a function of the actual number of days since mold insertion (Fig. 6). Data are shown separately for the 2008 (Fig. 6a-c) and 2010 (Fig. 6d-f) cohorts, because the 2008 cohort constituted the initial exploratory study and generally involved longer mold exposure, whereas the 2010 cohort was a more controlled follow-up study with mostly externally recruited participants and a shorter mold exposure (see Methods).

**Figure 6.**
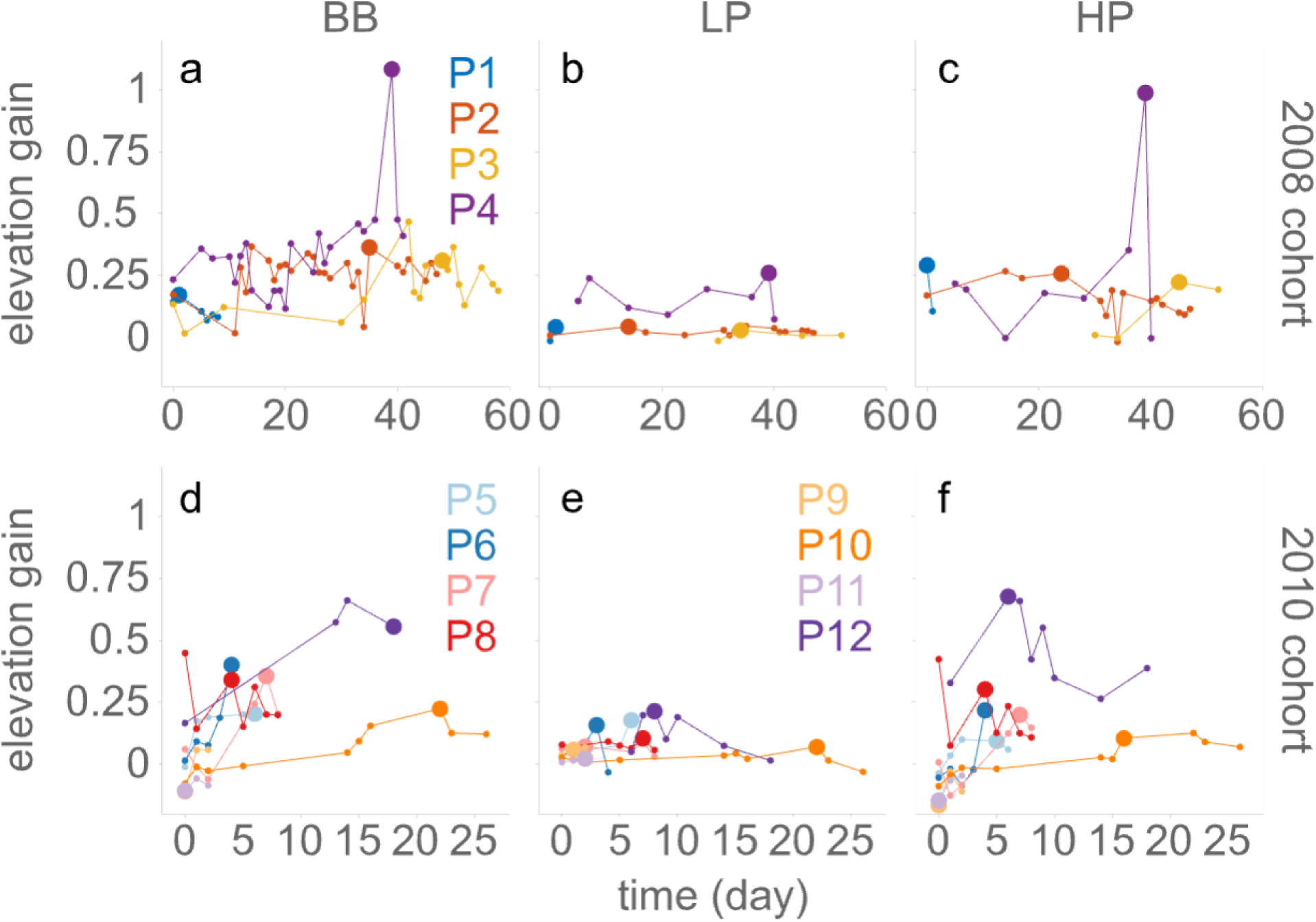
Time course of elevation gain adaptation. Elevation localization gains are plotted as a function of days since mold insertion for broadband (BB; a, d), low-pass (LP; b, e), and high-pass (HP; c, f) sounds. Data are shown separately for the 2008 cohort (a-c) and 2010 cohort (d-f). Each line represents data from one participant. The large, filled circle for each participant indicates the session with the highest target-response correlation for that participant and sound condition; these sessions were used as the maximal-adaptation measurements in Fig. 7. The gain trajectories show substantial inter-individual variability, session-to-session fluctuations, including in some cases decreases after peak performance.

The gain trajectories reveal substantial inter-individual variability and session-to-session fluctuations. For broadband sounds, several participants showed gradual increases in elevation gain, although this improvement was not always monotonic. High-pass sounds showed a similar pattern, whereas low-pass elevation gains generally remained close to zero. The larger markers indicate the session with the highest target-response correlation for each participant and sound condition, which was used for the maximal-adaptation analysis in Fig. 7. In several cases, performance declined after this session, e.g., P4, indicating that maximal observed performance should not be interpreted as a stable endpoint or asymptotic level of adaptation. Thus, the individual time courses do not support a simple monotonic relationship between exposure duration and the extent of adaptation. Rather, the amount of recovery depended strongly on participant, session, and stimulus condition, indicating that exposure duration alone did not explain the variability in performance.

**Figure 7.**
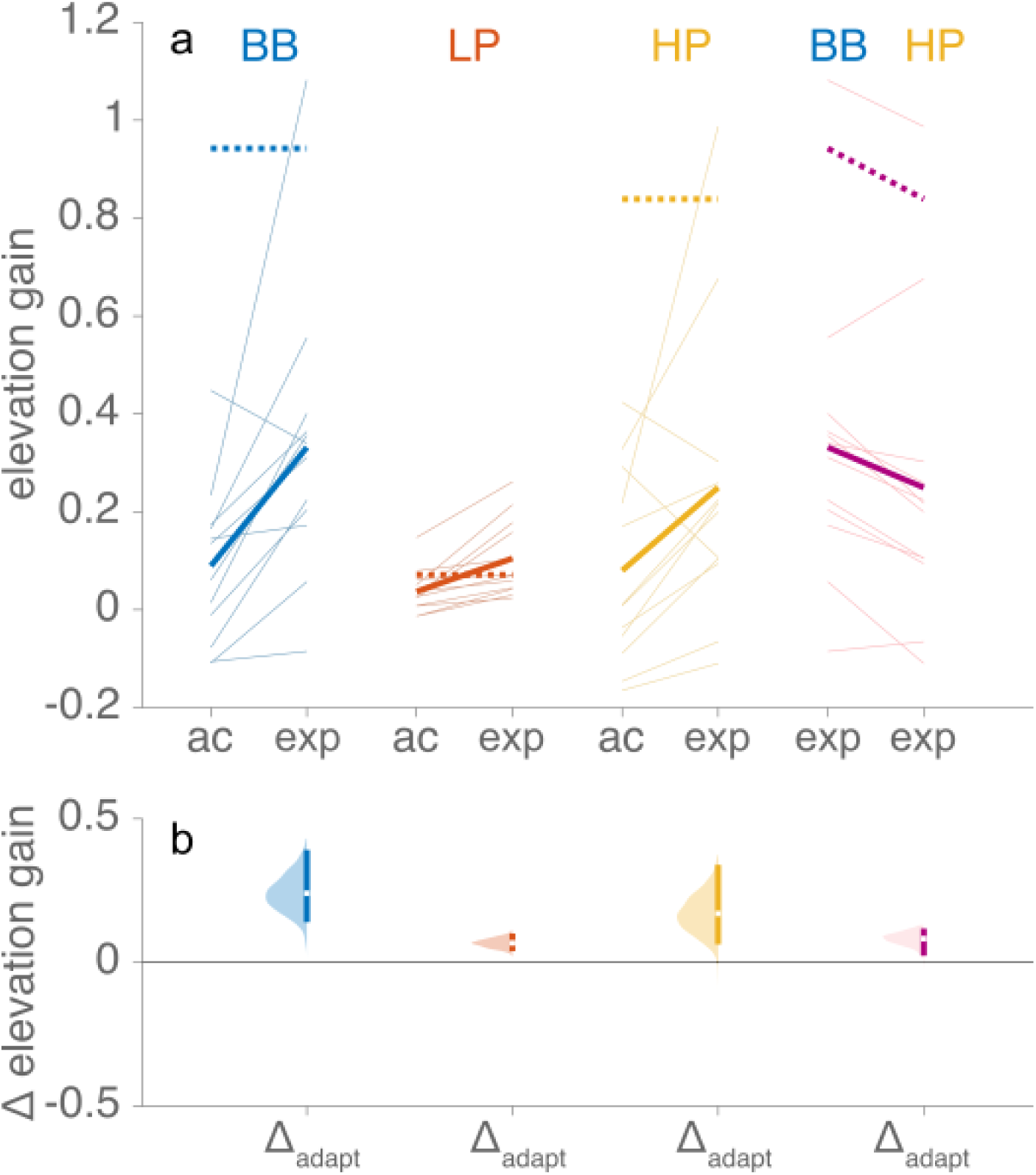
Adaptation of human sound localization performance to barn-owl molds. (a) Elevation localization gains measured with acutely worn barn-owl molds (“ac”) and after extended mold exposure (“exp”) for broadband (BB; blue), low-pass (LP; red), and high-pass (HP; yellow) sounds. Thin lines show individual participants; thick solid lines show group means for the with-mold condition, and dashed lines indicate baseline performance without molds. Purple symbols denote the direct comparison between broadband and high-pass conditions after extended use. (b) Paired mean differences in elevation gain between pre- and post-adaptation conditions. Error bars represent 95% confidence intervals, with the white marker indicating the mean difference. Colored patches depict the bootstrap sampling distributions of the differences.

### Adaptation to Molds

We next quantified the extent to which participants adapted to the altered spatial cues introduced by the barn owl molds after prolonged exposure. To that end, we compared localization performance immediately after mold insertion with the performance after extended use. For each participant, we selected the post-adaptation session with the highest elevation correlation. As before, we analyzed localization gains separately for elevation (Fig. 7) and azimuth (not shown), and for broadband (BB; blue), high-pass (HP; red), and low-pass (LP; yellow) sounds.

Elevation localization showed a modest but reliable improvement over the course of mold exposure, consistent with partial adaptation. For broadband sounds, elevation gain increased significantly from the acute phase to the session with the highest correlation during the exposure period (Δ_BB_ = 0.24 [0.15, 0.41]; *p* = 0.003; Fig. 7b), indicating substantial recovery relative to the immediate post-mold deficit. Localization of high-pass sounds likewise showed a significant gain increase over time (Δ_HP_ = 0.17 [0.06, 0.33]; *p* = 0.02; Fig. 7b), although the magnitude of recovery was smaller than in the broadband condition. Gain improvement was smallest for low-pass sounds (ΔLP = 0.07 [0.04, 0.10]; p ≪ 0.01; Fig. 7b), as expected because elevation localization for low-pass sounds was poor throughout the experiment.

Individual changes were variable, but the direction of change was generally consistent. Broadband elevation gains increased from the acute to the maximal-adaptation session in 11 of 12 datasets, whereas high-pass gains increased in 10 of 12 datasets. Thus, the group-level improvement was not driven by a single participant, although the magnitude of recovery differed substantially across listeners. The qualitative pattern of modest and variable elevation recovery, with the strongest improvement for broadband sounds, was also unchanged across robustness checks that excluded authors, accounted for differences in session timing, removed the highest-gain observation, or restricted the analysis to the 2010 cohort.

Adaptation remained incomplete and differed across sound types. Direct comparisons at the end of the exposure period revealed that elevation gains for broadband sounds exceeded those for high-pass sounds (Δ_BB-HP_ = 0.08 [0.03, 0.12]; *p* = 0.01; Fig. 7b), indicating a selective benefit for stimuli containing both low- and high-frequency components. This pattern suggests that successful elevation adaptation was facilitated by access to the full set of spatial cues available in broadband sounds, whereas recovery for high-pass stimuli alone was more limited. Because broadband and high-pass sounds both contained high-frequency spectral information, the additional recovery for broadband sounds points to a contribution of low-frequency spatial cues rather than to spectral-cue learning alone. Overall, these results demonstrate gradual but partial elevation adaptation, with clear evidence that the extent of recovery depended on stimulus cue content.

We also examined whether the primary mapping between target and response azimuth changed with prolonged mold exposure. Azimuth localization gains remained largely unchanged after mold insertion for broadband, low-pass, and high-pass sounds (Δ_BB_ = −0.06 [−0.18, 0.03]; *p* = 0.31; Δ_LP_ = 0.07 [−0.01, 0.14]; *p* = 0.10; Δ_HP_ = 0.00 [−0.09, 0.08]; *p* = 0.94).

Thus, the molds did not reliably affect horizontal localization accuracy. Since the molds acutely introduced a weak elevation-dependent modulation of azimuth localization responses (Fig. 5c,f), we tested whether this cross-dimensional effect was itself subject to recalibration by quantifying the dependence of azimuth response on target elevation across conditions. This elevation-dependence did not differ between the no-mold and acute-mold conditions for broadband, low-pass, or high-pass sounds (Δ_BB_ = 0.03 [−0.03, 0.09]; *p* = 0.42; Δ_LP_ = 0.02 [−0.06, 0.05]; *p* = 0.67; Δ_HP_ = 0.05 [−0.08, 0.16]; *p* = 0.46). Moreover, the relative difference between broadband and high-pass sounds was unchanged (Δ = 0.02 [−0.07, 0.08]; *p* = 0.61). Thus, both acutely and throughout adaptation, azimuth responses retained a small but stable elevation-dependent component that did not undergo recalibration.

## Discussion

### Summary of main findings

In this study, we examined the adaptability of human sound localization when listeners were confronted with radically altered spatial cues produced by asymmetric molds. Acute exposure to the molds disrupted the normal cue structure: pinna-based spectral cues were largely eliminated, high-frequency interaural level differences (ILDs) were altered, and low-frequency interaural time differences (ITDs) remained largely intact (Fig. 3). Consequently, elevation localization for broadband and high-pass sounds deteriorated severely, whereas low-pass elevation localization, already poor at baseline, remained largely unchanged. Horizontal localization was largely preserved, although the altered ILDs introduced systematic elevation-dependent biases in azimuth responses (Figs. 4 and 5). With prolonged exposure, listeners showed limited recovery of elevation localization, but this adaptation was slow, variable across participants, and fluctuating across sessions (Figs. 4, 6, 7). Importantly, recovery was strongest for broadband sounds, indicating that access to both low-frequency ITDs and high-frequency level cues facilitated adaptation. Together, these findings show that the human auditory system can adapt, albeit weakly and inefficiently, to cross-dimensional changes in spatial cues. The selective broadband advantage suggests that low-frequency ITDs helped listeners resolve the ambiguity of the altered ILD structure, allowing binaural cues normally associated with azimuth to contribute to elevation recovery.

### Acute cue perturbations reveal obligatory mapping of ILDs to azimuth

Sound localization is a computational process that relies on the interpretation and integration of multiple auditory spatial cues, including ITDs, ILDs, and monaural spectral cues (Strutt, 1907; Wightman and Kistler, 1989; Middlebrooks and Green, 1991; Blauert, 1997; Grothe et al., 2010; Joris and Van der Heijden, 2019; Yin et al., 2019). The asymmetric molds used here successfully generated acoustic transformations while disrupting normal human spatial cues (Fig. 3), and this manipulation immediately affected sound-localization behavior (Figs. 4 and 5). Such acute effects are consistent with more than a century of research demonstrating that abrupt alterations of spatial cues, through ear plugging, pinna molds, acoustic tubes, or virtual acoustics, produce predictable and often severe localization errors (e.g. Strutt, 1907; Wightman and Kistler, 1989; Hofman et al., 1998; Carlile, 2014; Kumpik and King, 2019). Each of these manipulations alters the spatial cues in a distinct way, thereby changing the mapping between cue values and perceived sound location. In the present study, the molds disrupted monaural spectral cues and introduced ILDs that varied systematically with both azimuth and elevation, while leaving ITDs unchanged. As a consequence, azimuth localization of high-pass sounds became dependent on sound-source elevation. Because the brain has no immediate access to the generative structure of the altered cues, initially, ILDs remain mapped onto azimuth alone, resulting in systematic cross-dimensional interference.

Interestingly, localization of broadband sounds showed a much weaker elevation dependence of azimuth responses and instead resembled localization of low-pass sounds (Fig. 5c). This pattern suggests that, for broadband sounds, low-frequency ITDs and high-frequency ILDs were combined in a weighted fashion, with ITDs exerting the dominant influence on azimuth judgments. Such weighted cue integration is consistent with classic ITD–ILD trading relations (e.g., Blauert, 1997; Macpherson and Middlebrooks, 2002; Heller and Richards, 2010; Stecker, 2010; Ahrens et al., 2020). ITDs provide a robust cue to horizontal sound location: within the low-frequency range in which they are most useful, they are only weakly frequency-dependent and are little affected by changes or obstructions in the external ear canal. By contrast, ILDs are strongly frequency-dependent and are more susceptible to changes in the external ear. The molds substantially altered the high-frequency ILD structure, such that ILDs no longer varied only with azimuth, as in normal listening, but also systematically with elevation. This introduced an ambiguity in the interpretation of ILDs: the same or similar ILD values could now reflect differences in azimuth, elevation, or both. This likely reduced the effective reliability of ILDs as a cue to azimuth and led the auditory system to down-weight ILDs relative to ITDs, in accordance with Bayesian principles of reliability-based cue integration (Ernst and Banks, 2002; Battaglia et al., 2003; Körding and Wolpert, 2004). Thus, the acute effects of the molds suggest that, immediately after the acoustic input was altered, listeners continued to rely primarily on their established (“prior”) cue-to-space mapping, with robust low-frequency ITDs dominating azimuth judgments even when the high-frequency ILD structure no longer conformed to the normal mapping.

### Acute pinna cue perturbations reveal their importance in sound elevation localization

The degradation of spectral pinna cues yielded clear and immediate deficits in elevation localization (Fig. 5a). This result is consistent with extensive previous work showing that high-frequency, monaural spectral features introduced by the pinna are the dominant cues for human sound-source elevation localization (Hebrank and Wright, 1974; Musicant and Butler, 1984; Middlebrooks, 1992; Hofman and Van Opstal, 1998; Langendijk and Bronkhorst, 2002). When these cues are removed or distorted by ear molds, earplugs, or spectral smoothing, elevation judgments typically collapse toward chance, even when binaural cues remain intact (Hofman et al., 1998; Macpherson and Sabin, 2013; Carlile, 2014). The present findings confirm that, in the absence of reliable spectral structure, neither ILDs nor ITDs alone provide sufficient information for accurate elevation localization under normal cue mappings, emphasizing the critical role of pinna-based spectral cues for normal vertical sound localization in humans.

### Broadband advantage supports adaptation to the altered cue structure

Despite the severe disruption of normal elevation cues, elevation gains improved over days to weeks, but recovery remained incomplete and fluctuated across sessions (Figs. 6, 7), with performance generally remaining below baseline even when individual participants reached relatively high gains at particular time points (Hofman et al., 1998; Van Wanrooij and Van Opstal, 2005). Crucially, this improvement was strongest for broadband sounds, whereas high-pass sounds showed substantially less recovery. This pattern is informative for interpreting the underlying mechanism. Recovery for high-pass sounds may reflect learning of residual or altered high-frequency spectral cues, altered high-frequency ILDs, or a combination of both. However, the additional recovery for broadband sounds over high-pass sounds cannot be explained by access to additional high-frequency spectral information, because both stimulus types contained such cues. The broadband advantage therefore suggests that listeners benefited from the combination of low-frequency ITDs and high-frequency mold-induced level cues, consistent with partial use of the altered binaural cue structure for elevation.

The adaptation data also showed substantial individual variability. Some participants showed little or no recovery, whereas others reached higher elevation gains during the exposure period. However, this variability is difficult to interpret mechanistically, because participants differed in exposure duration, number of test sessions, mold geometry, and possibly in daily acoustic experience. In addition, elevation gains fluctuated across sessions, so a single “adapted” value may not reflect stable performance. We therefore do not interpret individual differences as evidence for distinct learning strategies, but as an indication that adaptation to these altered cues was weak, unstable, and dependent on the specific conditions experienced by each listener.

The persistence of the small cross-dimensional bias, in which response azimuth depended on stimulus elevation (Figs. 4g,i, 5c), adds another constraint on the interpretation of adaptation. It indicates that the altered high-frequency level cues were not fully reassigned from azimuth to elevation, and that the original ILD–azimuth mapping remained influential. Thus, the present results should not be interpreted as complete remapping of ILDs onto elevation.

Because the molds altered several acoustic cues simultaneously, the present experiment does not isolate pure ILD-cued elevation learning. Residual or altered high-frequency monaural spectral cues may have contributed to recovery, especially for high-pass sounds. The molds also changed the spectra differently at the two ears, potentially creating interaural spectral differences that were not present in normal hearing. Such cues would be difficult to interpret unambiguously, because they depend not only on source direction but also on the spectrum of the sound source itself. Thus, unlike normal pinna-based elevation cues, they would constitute an ill-posed and source-dependent cue to auditory space, and hence inherently ambiguous. However, spectral-cue learning alone does not explain the additional broadband advantage, because broadband and high-pass sounds contained identical high-frequency spectral information.

Likewise, a contribution from low-frequency head- or torso-related cues cannot be fully excluded. Low-pass elevation gains showed a small average increase over the exposure period, suggesting that listeners may have extracted some elevation-related information even from the low-frequency stimulus band. However, this effect was much smaller than the recovery observed for broadband and high-pass sounds, and the within-participant pattern did not account for the additional broadband advantage over high-pass sounds. Moreover, low-frequency head- or torso-related cues are likely to be weak and posture-dependent. In everyday listening, head-on-torso posture varies independently of sound-source elevation, making such cues unreliable for driving adaptation from incidental feedback. We therefore interpret the broadband advantage as most consistent with the joint availability of low-frequency ITDs and altered high-frequency level cues, while acknowledging that low-frequency monaural cues may have contributed modestly to performance.

### Cue integration and possible neural locus of adaptation

Sound localization is not a direct readout of physical space, but depends on the extraction, integration, and interpretation of acoustic cues. In normal human hearing, ITDs and ILDs provide reliable information about azimuth, whereas high-frequency monaural spectral cues generated by the pinnae provide the dominant information about elevation.

Other cues, such as overall monaural intensity, may also influence localization (especially in impaired hearing; Van Wanrooij and Van Opstal, 2004), but generally have a less specific, often ambiguous, relationship to spatial position. Thus, the auditory system must extract spatial information from the spectrotemporal signals reaching the two ears, combine the resulting cue information, and map the resulting neural activity onto perceived sound location and orienting behavior.

The elementary spatial cues are first extracted in specialized subcortical circuits. ITDs and ILDs are processed in binaural brainstem pathways involving the medial and lateral superior olives, whereas monaural spectral information is transformed through cochlear-nucleus pathways, including the dorsal cochlear nucleus (Young et al., 1997; Grothe et al., 2010; Yin et al., 2019; Joris and Van der Heijden, 2019; Tollin, 2003). Spatial information is then progressively integrated in the midbrain and forebrain. Neurons in the inferior colliculus receive convergent information related to sound level, ITDs, ILDs, and spectral cues, and even eye orientation (Groh et al., 2001; Zwiers et al., 2004), and may therefore provide a distributed population code from which the sound location for eye-head orienting can be inferred (Zwiers et al., 2004; Chase and Young, 2008; Slee and Young, 2011, 2014; Yao et al., 2015; Van den Wildenberg and Bremen, 2024). However, unlike in the barn owl, a stable point-to-point auditory map of space has not been identified in the mammalian auditory pathway (Zwiers et al., 2004; Middlebrooks, 2021). This suggests that sound location in mammals is likely represented by distributed neural activity that must be decoded for perception and action.

A useful framework for the decoding of inferior colliculus spatial information was proposed by Zwiers et al. (2004). In their model, a tonotopic population of IC neurons, with distributed and apparently non-map-like activity, weakly modulated by eye position, projected to a spatially organized eye-centered motor map in the superior colliculus. By training the weights of the IC–SC projection, the network learned to transform distributed auditory activity into an eye-centered Gaussian representation of eye-motor error in the SC. Thus, in mammals, a spatially organized orienting signal need not require a topographic auditory space map at the level of the IC itself; it could emerge through learned weighting of distributed auditory activity onto a downstream sensorimotor map. Barn owl studies provide an important experimental counterpart to this idea. In the owl, auditory spatial information is represented in the external nucleus of the inferior colliculus (ICx), which projects to the optic tectum, the avian homologue of the superior colliculus. During prism adaptation, altered visual experience shifts auditory orienting behavior and the auditory space map, and this plasticity has been linked to changes in the pathway from the central nucleus of the inferior colliculus (ICc) to ICx and onward to the optic tectum (Brainard and Knudsen, 1993; Knudsen, 1983, 1999, 2002; Knudsen et al., 2000; Gold and Knudsen, 2000; Gutfreund et al., 2002; Peña and DeBello, 2010). Thus, whereas Zwiers et al. provide a computational account of how distributed auditory activity could be mapped onto a superior-colliculus motor representation, the barn owl work demonstrates experimentally that auditory-to-orienting mappings in the midbrain can be modified by experience. Note, however, that the barn owl has an extremely limited oculomotor range (<2 deg), making its head-centered auditory space map in the ICx ready for direct readout by the optic tectum for head orienting. In contrast, cats and primates use eye movements, which leads to dynamic misalignments of head-centered auditory space and eye-centered visual space. This necessitates a radically different neural mapping strategy than in the barn owl.

This distinction is important for interpreting the present results. Adaptation to the asymmetric molds need not require a change in early feature extraction. ITD-, ILD-, and spectral-cue-sensitive circuits may continue to encode the altered acoustic input according to their normal sensitivities. Instead, adaptation could occur by changing the downstream weighting or interpretation of the resulting cue-related population activity. In this view, the molds altered the pattern of activity entering integrative auditory stages, and experience gradually modified how that activity was mapped onto spatial judgments or orienting responses. The selective broadband advantage is consistent with such a cue-weighting or mapping account. With the molds in place, high-frequency level cues depended on both azimuth and elevation, whereas low-frequency ITDs remained a relatively stable cue to azimuth. For broadband sounds, listeners could therefore use ITDs to help resolve the azimuthal component of the altered ILD structure, allowing part of the high-frequency level information to contribute to elevation judgments. For high-pass sounds, this stabilizing low-frequency ITD information was not available, making the altered high-frequency cue structure more difficult to interpret. Thus, the same altered cue-related activity may have been weighted or decoded differently at later auditory, multisensory, or orienting stages, rather than requiring ILD-sensitive brainstem neurons to change their basic tuning.

The present data do not allow us to determine whether this adaptation occurred at a perceptual level, a sensorimotor mapping level, or both. However, we consider a purely decisional (cognitive) strategy unlikely. Participants were instructed to orient to the perceived sound location, were not told the acoustic structure of the manipulation, and received no trial-by-trial feedback during laboratory testing. Thus, they had little opportunity to learn an explicit response rule in the experimental setting. Moreover, the altered cue structure was not reducible to a simple rule such as mapping perceived azimuth onto elevation: the mold-induced ILDs varied with both azimuth and elevation, while ITDs remained informative about azimuth. A simple strategic rule would therefore be expected to introduce systematic errors in other response dimensions, rather than selectively improving broadband elevation localization. Finally, participants did not generally report a strong, explicit awareness of a systematic change in elevation perception during everyday listening, apart from occasional mislocalizations. We therefore interpret the recovery as behavioral adaptation to the altered spatial cue structure, most plausibly involving changes in cue weighting or cue-to-space mapping, rather than as an explicit decisional strategy.

### Why adaptation is limited: weak and ambiguous cues

Adaptation to altered auditory spatial cues has been demonstrated in many previous studies, but its rate and extent depend strongly on the nature of the cue perturbation (Van Wanrooij and Van Opstal, 2005; Carlile, 2014; Mendonça, 2014; Kumpik and King, 2019). When spatial cues are modified within an existing cue-to-space mapping, such as by unilateral ear plugging or pinna molds that preserve a consistent mapping between spectral features and elevation, humans can show substantial and sometimes rapid recalibration (Hofman et al., 1998; Van Wanrooij and Van Opstal, 2005; Kumpik et al., 2010; Carlile, 2014; Kumpik and King, 2019). In contrast, more extreme manipulations that disrupt the internal consistency of spatial cues, such as binaural input reversal, typically result in very slow adaptation or a lack thereof (Hofman et al., 2002; King et al., 2011). The limited adaptation observed in the present study is consistent with the latter studies.

The magnitude of recovery should be interpreted cautiously. Even for broadband sounds, where adaptation was strongest, elevation gains remained well below baseline. Moreover, these values represent the maximal observed recovery within the available exposure period, not necessarily final or asymptotic performance. Adaptation fluctuated across days, and the experiment ended before a plateau could be established. Thus, longer exposure might have yielded further improvement, although the present data suggest that any such learning would be slow and variable. The observed gains may have provided coarse elevation information but were insufficient for functional localization.

A key constraint on adaptation in our experiment was the altered relationship between spatial cues. The asymmetric molds introduced ILDs that varied with both azimuth and elevation, creating a cue mixture that does not exist in normal human hearing (Fig. 1b; Wightman and Kistler, 1989; Middlebrooks and Green, 1991). Successful adaptation would therefore have required more than recalibrating an existing cue-to-space mapping. A more complete adaptation would likely have required listeners to reinterpret ILDs while taking their azimuth dependence into account, so that binaural cues could provide more independent information about elevation. The persistence of elevation-dependent azimuth biases suggests that this remapping was incomplete.

This limitation may partly reflect the relatively weak and imbalanced elevation information carried by the mold-induced ILDs (Fig. 3b), especially when compared with the pronounced elevation-dependent ILDs available to barn owls (Fig. 1e). As illustrated for participant 2, elevation-dependent level changes were evident primarily for the left ear, whereas corresponding effects at the right ear were weak or absent. The resulting ILD pattern may therefore not have constituted a robust binaural cue that could be interpreted consistently as signaling elevation. Instead, listeners may have been confronted with an ear-specific level structure that was poorly matched to existing binaural comparison mechanisms. Stronger, more symmetric, or more systematic elevation-dependent ILDs might therefore support more robust adaptation.

The broadband advantage is consistent with this interpretation: low-frequency ITDs may have provided a stable azimuth reference that helped listeners interpret the altered high-frequency level structure. However, because the mold-induced ILDs still varied with both azimuth and elevation, disentangling these contributions likely remained difficult and limited the extent to which ILDs could be repurposed for elevation localization. Thus, the improvement in elevation gain should be interpreted as partial behavioral adaptation to an altered cue structure, rather than as evidence for complete reassignment of ILDs from azimuth to elevation. Future work using virtual-acoustic or wearable real-time cue transformations could impose more systematic and stronger spatial cue manipulations than the currently used physical molds, although maintaining such transformations during natural everyday listening remains technically challenging.

### Why adaptation is limited: imprecise error signals

A further constraint on learning may arise from a relationship between localization accuracy and precision. Previous studies have shown that localization gain and response variability are linked, particularly under conditions of increased sensory uncertainty (Ege et al., 2018; Ege et al., 2026; Garcia et al., 2017; Bruns et al., 2024). In a simple inference framework, reducing response gain can reduce response variability when sensory uncertainty is high, because responses are compressed toward the centre. Such a strategy may minimize variable errors, but it also creates a challenge for adaptation. With the molds, the sensory evidence for elevation was weak and ambiguous. Increasing gain during learning would therefore make responses more dependent on this uncertain sensory evidence, potentially increasing trial-by-trial errors even if average accuracy improved. Natural sensorimotor feedback may then provide inconsistent teaching signals: a partially learned mapping could appear useful on some occasions but misleading on others. This may contribute to the slow, fluctuating, and incomplete adaptation observed here. We explored response precision in the present data by estimating the residual variability of the stimulus-response fits, but these estimates were too noisy to draw strong conclusions about the accuracy–precision relationship or its time course during adaptation. Future studies with more trials per session could test whether precision limits the learning of altered spatial cue mappings.

### Limitations

Several limitations of the present study should be acknowledged, most of which arise from the practical burden of the experimental paradigm. The sample size was modest, and for many participants the duration of continuous mold use was limited. This is important because previous studies show that adaptation to altered external-ear cues can unfold over very different time scales: some listeners improve within days, whereas others require weeks or even months, depending on the cue perturbation, feedback, and individual listener (Hofman et al., 1998; Van Wanrooij and Van Opstal, 2005; Carlile et al., 2014; for review see Carlile, 2014). Thus, for some participants in the present study, the exposure period may have been too short to determine whether further adaptation would have occurred. Extending the exposure period, however, was constrained by the practical burden of the paradigm. Wearing asymmetric ear molds throughout daily life is physically uncomfortable and socially conspicuous, and repeated laboratory testing over days to weeks is time-consuming and fatiguing. Continuing the experiment until each participant reached a stable asymptote was therefore not feasible. These factors constrained both recruitment and exposure duration and therefore reflect unavoidable design trade-offs rather than experimental oversights.

A second limitation is that the molds were manually fabricated for each listener, so the acoustic transformations were unlikely to be identical across participants. Such variability may have contributed to the heterogeneous and fluctuating adaptation observed here. This interpretation is consistent with earlier mold studies, in which learning could be slow and variable when cue changes depended on individual ear morphology (Hofman et al., 1998; Van Wanrooij and Van Opstal, 2005). By contrast, more controlled manipulations, such as standardized molds or virtual-acoustic transformations, can impose more systematic cue changes and may therefore support clearer or faster learning (Zahorik et al., 2006; Majdak et al., 2013; Carlile et al., 2014).

Finally, participants received no explicit training or performance feedback during testing. This choice was intentional: by withholding feedback, we aimed to assess spontaneous adaptation driven by natural sensorimotor experience in everyday environments, rather than task-specific or setup-dependent learning. Although explicit training and feedback can accelerate and enhance adaptation to altered spectral cues, such approaches may also promote strategies tied to specific stimuli or loudspeaker configurations (Majdak et al., 2013; Mendonça, 2014; Keating et al., 2016; Kumpik et al., 2019; Arras et al., 2022; Kacelnik et al., 2002).

### Potential implications for hearing rehabilitation

The present findings may inform future cue-shaping approaches for individuals with hearing impairment, particularly those with limited access to high-frequency spectral cues (Ludwig et al., 2021; Zheng et al., 2022). Many common forms of hearing loss, as well as hearing aids and cochlear implants, degrade or eliminate the fine spectral structure generated by the pinnae, thereby compromising elevation localization (Otte et al., 2013; Thakkar and Goupell, 2014). Our results suggest that the auditory system may retain some capacity to exploit altered cue structures when multiple binaural cues are jointly available. However, the modest and fluctuating recovery observed here also shows that such relearning is strongly constrained when binaural cues are weak, ambiguous, or confounded by natural acoustics.

Listeners with hearing impairment differ from the normal-hearing listeners tested here because their acoustic input is shaped by hearing devices. In principle, hearing aids or cochlear implants could impose stronger and more systematic cue mappings than those produced by physical molds, for example by preserving ITDs for azimuth while using ILDs or other level-based transformations to support elevation. From this perspective, the limited adaptation observed here does not preclude rehabilitative benefit but highlights the importance of consistent cue structure and training. Device-mediated cue shaping, particularly when combined with explicit feedback or training (Keating et al., 2016; Kumpik et al., 2019; Arras et al., 2022), may allow listeners to exploit binaural cues for vertical spatial perception in ways that are not feasible through natural acoustics alone.

## Conclusion

By confronting human listeners with asymmetric outer-ear acoustics, we tested the limits of auditory spatial plasticity under extreme cue perturbations. The altered cues immediately disrupted elevation localization and induced cross-dimensional biases, revealing the persistence of established cue-to-space mappings. With prolonged exposure, listeners showed a modest but reliable improvement in elevation localization, which was strongest for broadband sounds. This pattern argues against an explanation of the broadband advantage based on relearning high-frequency spectral pinna cues alone, because broadband and high-pass sounds contained similar high-frequency information. Instead, the additional improvement for broadband sounds suggests that listeners benefited from the joint availability of low-frequency ITDs and altered high-frequency level cues, with ITDs helping to disambiguate elevation-dependent ILDs. Thus, the results indicate partial behavioral adaptation to an altered spatial cue structure, while also revealing strong constraints on the ability of the adult human auditory system to remap spatial dimensions.

## Conflict of Interest

The authors declare no competing interests.

## Author contributions

MW, RW designed the research; MW performed the experiments; MW, PB analyzed the data; MW, PB drafted the manuscript; JO, MW, PB, RW wrote the paper; all approved the final version of the manuscript.

## Data availability

The data supporting the findings of this study are available from the Radboud Data Repository at https://doi.org/10.34973/fktt-7246.

## Funding

This work was supported by the European Commission, Marie Curie Early-Stage Training Fellowship, Sixth Framework Programme (MEST-CT-2004-007825, PB), Vici Grant ALW 865.05.003 within Earth and Life Sciences of The Netherlands Organization for Scientific Research (JO, MW, RW), and Radboud University (JO, MW).

## Acknowledgements

We thank Sam Bruijs and Bart Gips for their help in the initial phase of this project. We thank Günter Windau, Ruurd Lof, Dick Heeren, Hans Kleijnen and Stijn Martens for valuable technical assistance and Arno Engels and Jaap Nieboer for engineering and building the auditory hoop. We also thank the five anonymous reviewers for their careful and constructive feedback, which substantially improved the manuscript.

## Notes

https://doi.org/10.34973/fktt-7246

### Competing Interest Statement

The authors have declared no competing interest.

### Summary of Updates

The manuscript was substantially revised to improve clarity, rigor, and transparency; to moderate the interpretation of adaptation to altered binaural cues; to better address alternative explanations and limitations; and to clarify the acoustic cue transformations, individual variability, and incomplete nature of the observed learning.

